# A computational architecture incorporating shallow brain networks: integrating parallel cortical and subcortical processing

**DOI:** 10.1101/2025.02.24.639932

**Authors:** Kwangjun Lee, Lorenzo Baracco, Cyriel M. A. Pennartz, Mototaka Suzuki, Jorge F. Mejias

## Abstract

Artificial neural networks commonly have deep hierarchical structures that were originally inspired by the neuroanatomical evidence of cortico-cortical connectivity pattern found in the mammalian brain. Largely neglected in those models are non-hierarchical aspects of brain architecture, namely the subcortical pathways and the interactions between cortical and subcortical areas regardless of their hierarchical locations. Inspired by this principle, we present a computational model combining cortical hierarchical processing with subcortical pathways based on neuroanatomical evidence. We show the versatility of our model by implementing the cortical hierarchy in two alternative ways—a convolutional feedforward network and a predictive coding network. Both model variants can replicate behavioral observations in humans and monkeys on a perceptual context-dependent decision-making task. The model also reveals that subcortical structures lead decisions for easy trials while the more complex hierarchical network is necessary for the harder trials. Our results suggest that the parallel cortico-subcortical processing explored in the model represents a fundamental property that cannot be neglected in understanding the computational principles used by the brain.

**Significance:** Artificial intelligence and computational neuroscience models, particularly deep learning and predictive coding architectures, have been traditionally dominated by cortico-centric hierarchical frameworks. However, extensive neurobiological evidence suggests that cortical areas, regardless of their hierarchical classification, are deeply interconnected with subcortical structures in real brains. We propose here a computational framework demonstrating that parallel cortical-subcortical architectures can yield complex and more flexible computational capabilities, aligning with the observed behavior in the mammalian brain. Our model addresses key limitations in existing deep learning and predictive coding networks, offering a more biologically plausible and functionally advantageous alternative.

## Introduction

Artificial Intelligence (AI) is currently transforming nearly all aspects of our lives in an unprecedented way^1,2^ and in particular the way we think about how the brain works^1,3^. Today’s dominant AI models for visual perception are based on Deep Learning (DL) architectures—those possessing deep hierarchical structures that consist of a stack of layers, each of which contains a varying number of artificial neurons^1^. Historically, such DL architectures were inspired by neuroanatomical findings in the visual system of the primate neocortex a few decades ago^1,4^. Tracing studies found a pattern of connectivity between cortical areas^5,6^; according to the rules of axonal termination, cortical areas appear to have a deep hierarchical structure^5,6^.

However, as highlighted by recent work^7^, cortico-cortical connections forming this hierarchical structure are merely a subset of the total number of connections that cortical areas actually have. Connections neglected in DL models include, for example, local recurrent synapses^8,9^, dense recurrent long-range connections^10,11^ and, in particular, the connections from every cortical area—regardless of whether the cortical area is hierarchically high or low—to subcortical regions such as the brainstem, thalamus, and striatum^7^. Another set of connections underrepresented in today’s DL models is the collection of projections every cortical area receives directly from subcortical regions^7^. Currently, there is no computational model that integrates all these reciprocal connections between cortical and subcortical areas in a DL framework, let alone the interactions between the cortico-subcortical communication and the hierarchical processing system. The current AI models that depend solely on deep hierarchical structures may therefore be far from the computations performed in the brain and missing some of its fundamental computational principles.

To improve our understanding of the ways the brain processes sensory information to facilitate cognitive computations, we need to take the connections neglected in today’s DL models into consideration. Likewise, we must computationally explore the function of those connections, because anatomical evidence suggests that they play a fundamental role^7^. In the present work, we take the first step by examining one of such neglected anatomical features of the brain, namely, the subcortical pathways that process information in parallel with cortical pathways. We present a computational framework that combines a deep cortical hierarchy and shallow subcortical pathway, and it does so in two possible implementations: a feedforward network, and a predictive coding network. We replicate some of the characteristic behavior observed in human and nonhuman primates, such as rule-dependent distributions of reaction times (which cannot be accounted for by the hierarchical cortical processing alone) and provide predictions regarding the importance of cortical and subcortical structures as a function of the task difficulty.

## Results

Our goal in this study is to explore how a model incorporating deep hierarchical cortical and shallow subcortical systems may explain complex behavioral tasks which cannot rely on deep architectures alone. We decided to implement a prosaccade/antisaccade task^12^ similar to those used in decision-making experiments for macaques^13^, given its clear distinction between easy and hard trials. In this task (Fig. 1A), the macaque receives at the beginning of each trial a visual cue, which indicates whether the trial is prosaccade or antisaccade. For a prosaccade trial, two stimuli of different contrasts are simultaneously presented in the screen, and the macaque has to saccade towards the most salient one (i.e. the one with the highest contrast). Such purely luminance-driven response may be likely achieved via rudimentary reflexive responses which do not require advanced perception and cognition in the animal^14^. On the antisaccade trial, however, the monkey must suppress the urge to saccade towards the brighter stimulus and look instead at the one with lower contrast. This operation likely involves circuits in higher cortical areas in charge of inhibitory control and flexible behavior^15–19^.

**Figure 1.**
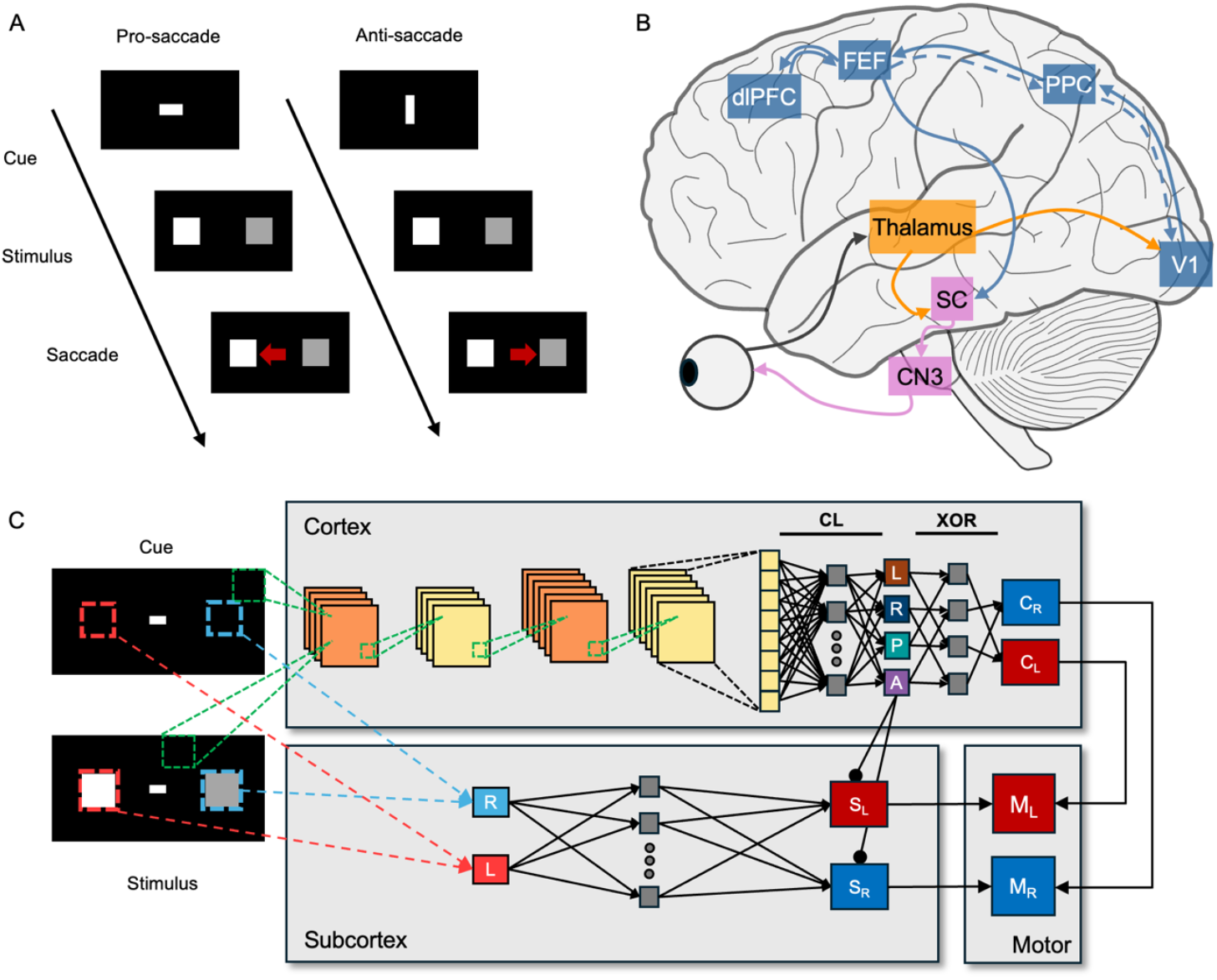
A feedforward model implementation of the shallow brain hypothesis. (A) The behavioral task adopted here. Each trial began with a horizontal or vertical bar cueing the model to perform a pro-or antisaccade. In prosaccade trials, the model made a saccade towards the brighter square (left), while in antisaccade trials, it made a saccade away from it (right). (B) The neuroanatomical pathways considered in the model. We consider a deep cortical hierarchy hypothetically involving the thalamus as well as cortical areas like V1, PPC, FEF, and dlPFC. The FEF is assumed to code target stimulus location and task rule (P or A) and send an effective inhibitory cortico-subcortical projection to the superior colliculus, SC. The oculomotor commands are generated by combining subcortical and cortical outputs. (C) The model architecture. The cortical processing pathway classified the cue indicating the task rule (pro, P, or anti, A) and later integrated it with the subsequent outcome of brightness comparison (left, L, or right, R) to generate a choice output (left or right choice; C_L_ or C_R_). CL indicates the classifier, output of which is compared to labels for the training via backpropagation. XOR indicates the hard-wired connectivity functioning as an XOR gate that integrates sequential classification results of the cue and brightness comparison. The subcortical processing pathway makes a quick decision with two subcortical neurons (R or L), the receptive fields of which cover the region where left or right square was located. Cortico-subcortical connections (A→S_L_ and A→S_R_) suppressed subcortical outputs during antisaccade trials. The final saccadic decision and RTs were read from motor neuron firing rates (*M*_*L*_ and *M*_*R*_), which combined outputs from cortical and subcortical pathways.

To better understand how these pro- and antisaccade trials can be processed by the brain, we identified cortical and subcortical neuroanatomical pathways involved in the perceptual and decision components of the task (Fig. 1B). We first considered that retinal ganglion cells (RGCs) send projections to not only the lateral geniculate nucleus in the thalamus that in turn projects to the primary visual cortex, but RGCs also directly project to superior colliculus (SC)^20^. In primates, SC directly projects to three cranial motor nuclei that control six eye muscles^21^. This subcortical pathway, which plays an important role in oculomotor control, is parallel to the well-studied dorsal cortical hierarchical pathway involving primary visual cortex (V1), posterior parietal cortex (PPC), and frontal eye field (FEF). Similar subcortical pathways exist also in other sensory modalities besides vision^22,23^, suggesting that such pathways may have a fundamental role in perceptual computations.

To explore how these two structural pathways cooperatively function in parallel, we took inspiration from our recently proposed shallow brain hypothesis^7^ and implemented a computational model accommodating hierarchical and shallow networks. As the LGN projects directly to the SC, we implemented a subcortical pathway that is significantly shorter compared to the cortical hierarchical pathway. To explore the versatility of our framework, we designed and implemented the cortical network in two alterative architectures: a convolutional feedforward network and a predictive coding network.

### Combining a convolutional feedforward cortical network with a shallow subcortical network

As a first implementation, we adopted a network with two subparts to model the cortical processing pathway, and a simple network for the subcortical pathway (cortex; Fig. 1C). The first cortical subpart, a convolutional feedforward pathway, made a four-way classification among two cues (pro-or antisaccade; P and A in Fig. 1C) and two brightness comparison outcomes (left square brighter than right or vice versa; L and R in Fig. 1C). The second cortical subpart, an implicit memory and decision-making pathway, saved the classification outcome during the cue presentation and later combined it with the subsequent outcome of brightness comparison to make cue-based choices. The subcortex acted as a simple two-layer multilayer perceptron (MLP) with a single hidden layer. The subcortical module consisted of two units (representing two neuronal populations), the receptive fields of which cover the two areas where a pair of brightness stimuli appeared (red and blue dotted lines on cue and stimulus screen; Fig. 1C). A cortico-subcortical projection (A→S_L_ and A→S_R_ in Fig 1C; FEF → SC in Fig. 1B) inhibited the subcortical outputs only during antisaccade trials^24^. The motor neurons (L_M_ and R_M_) combined outputs from the cortical (C_L_ and C_R_) and subcortical (S_L_ and S_R_) modules to make a final choice.

The feedforward model performed near 100% accuracy during both pro- and antisaccade trials. This classification performance was robust against noise, suggesting that the model had learned good representations of inputs and consistently performed above chance level (>75%; Fig. 2A).

**Figure 2.**
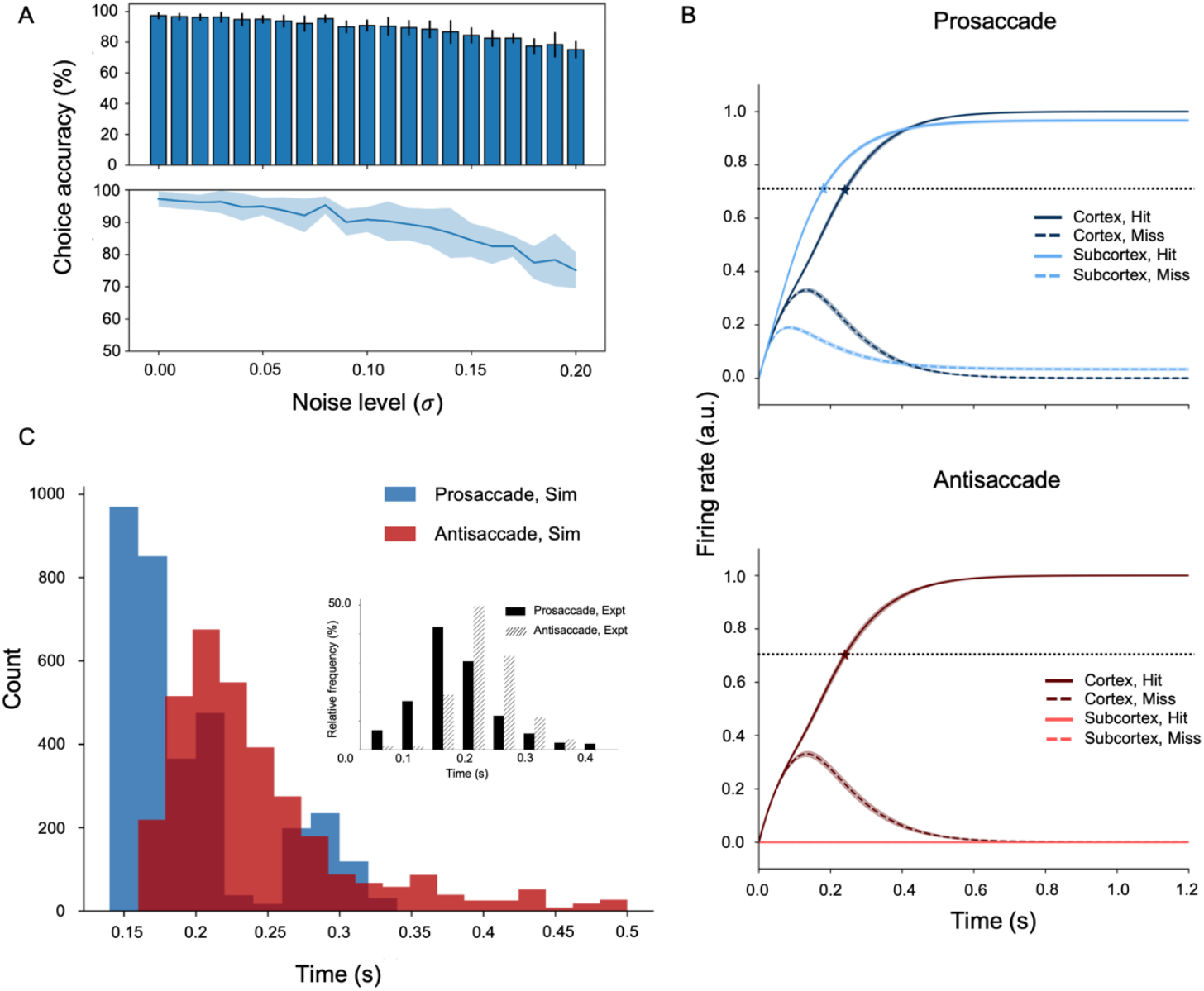
Model combining a feedforward cortical and subcortical network performing cue-based antisaccade. (A) Robustness against noise. The model consistently outperformed chance level across increasing levels of input noise. (B) Subcortical and cortical contributions to saccade decisions. During prosaccade trials, the subcortical module (solid light blue) rapidly drives the saccade decision. During antisaccade trials, cortico-subcortical inhibition (solid and dashed light red) silences the subcortex, allowing the cortex to guide the saccade. The horizontal dotted black line indicates a decision threshold. Stimulus onset is at 0.0 s. (C) Faster reaction times (RTs) during prosaccade (blue) than antisaccade (red) trials captured the RT pattern observed in monkey experiments (filled and hatched black)^13^: 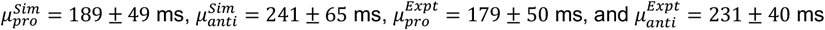

To compare our simulations with physiological data on antisaccade experiments in monkeys^13^, we computed reaction times (RTs) by recording the time at which either the left or right motor neuron (solid vs dashed line in blue and red hues; Fig. 2B) fired above a decision threshold (the dotted black line; Fig. 2B). Behavioral output (i.e., saccade direction) was determined by the first motor neuron to fire above the decision threshold. Interestingly, the simulated RT distribution showed similar patterns with the experimental result from an antisaccade task (Fig. 2C): not only was the mean RT faster for prosaccade than antisaccade trials (189 and 241 ms, respectively), but the actual values also fell within ranges of those obtained in the experiment (179 ± 50 ms and 231 ± 40 ms, respectively).

To analyze which of the two pathways, cortical or subcortical, contributes more to saccade decisions, we compared the mean firing rates of decision neurons in subcortical (*S*_*L*_ and *S*_*R*_) and cortical (*C*_*L*_ and *C*_*R*_) pathways separately. We found a clear distinction between correct and incorrect choices with minimal variance across trials (solid and dashed lines in blue and red hues; Fig. 2B). Subcortical outputs reached the firing threshold faster than cortical outputs during prosaccade trials (solid light vs dark blue lines in top panel; Fig. 2B). Meanwhile, the cortico-subcortical projections ensured that the subcortical contribution during antisaccade trials was minimal (solid and dashed lines in bright red near zero in bottom panel; Fig. 2B).

### Combining a predictive coding cortical network with a shallow subcortical network

Predictive coding (PC) networks can explain salient, biologically plausible features of brain function that fall beyond the scope of more traditional deep network structures, such as the ability of generating internal representations (or ‘world models*i*) of causes of sensory input^25–28^. They are also able to achieve a performance equivalent to deep architectures in some tasks without the need for error backpropagation for their training^29–31^. This motivated us to model the cortical hierarchy with a PC network (Fig. 3A) as an alternative for the deep network structure adopted in the previous section. A three-layer PC network was designed to model the cortical hierarchy, in parallel with a simplified version of recurrent network for perceptual decision-making^32^ to model the shallow subcortical pathway.

**Figure 3.**
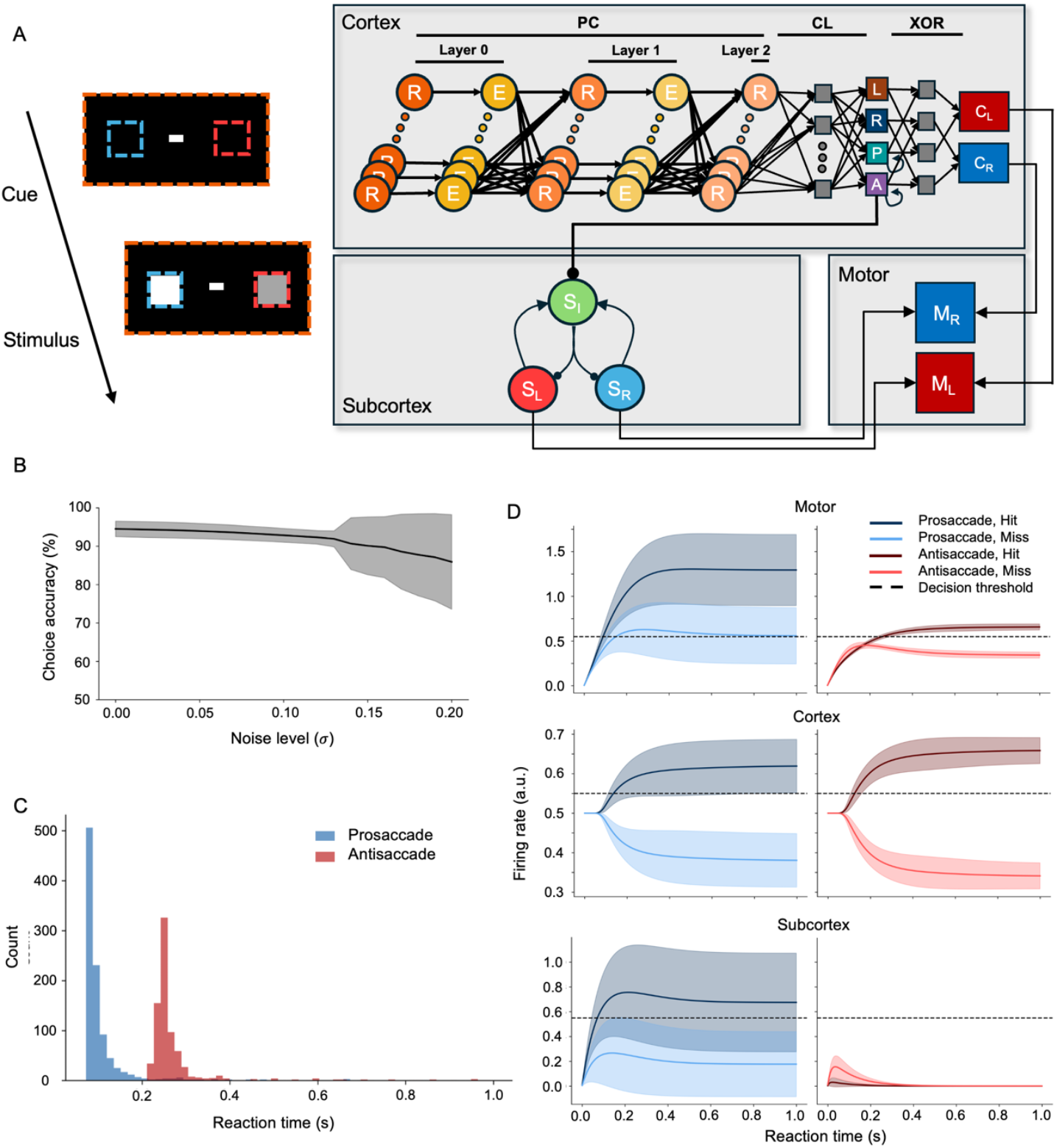
*M*odel *combining* a predictive coding and shallow-brain network. (A) Model architecture. The cortical pathway comprised three predictive coding layers (layers 0-2), roughly corresponding to V1, PPC, and FEF (Fig. 1B) and a linear classifier (CL) approximating the decision-making functionality of dlPFC. Sequential classification results of the cue and brightness comparison are integrated via a hard-wired XOR gate (XOR) to make cue-based decisions. The antisaccade cue classifier inhibits subcortical activity during antisaccade trials via a cortico-subcortical projection (A→S_I_). Motor neurons (M_L_ and M_R_) integrate cortical and subcortical outputs to make a saccade. (B) Robustness against noise. Task performance was consistently higher than chance level across the whole range of Gaussian noise levels added to inputs. The gray highlighted region indicates the standard deviation. (C) Simulated reaction time (RT) distributions show that saccades were made faster during prosaccade than antisaccade trials. (D) Subcortical and cortical dynamics are similar to the feedforward model (Fig. 2C). Mean firing rates of motor neurons and cortical and subcortical decision neurons during the saccade/antisaccade task. During prosaccade trials, the subcortex module drove the saccade decisions (solid dark blue > solid light blue). During antisaccade trials, the subcortex module was silenced by an inhibitory cortico-subcortical projection (solid and dashed light red; middle panel). In both cue conditions, the model made the correct saccade decisions (top panel).

The PC model was trained by a local learning rule^26^ to achieve performance equivalent to the feedforward network used in the previous section: robust classification performance against low and moderate levels of noise (Fig. 3B), delayed RTs during antisaccade trials (Fig. 3C), and inhibited subcortical responses during antisaccade trials (Fig. 3D).

The PC model reconstructed images in the input layer using predictive information from each level of the visual hierarchy (Fig. 4A), indicating the formation of appropriate hierarchical representations of visual inputs. To take advantage of richer representations of generative models compared to purely discriminative models (e.g., the feedforward CNN), we asked whether the PC model contains/codes information about task difficulty. Trials were first divided into two groups: those trials in which motor outputs were driven by the subcortex and other trials driven by the cortex. To determine whether the cortex or the subcortex drove the decision, we looked at the final output from each module to motor neurons (Fig. 4A) and selected the module that reached the decision threshold before the other module as a winner. Then, we sorted trials according to task difficulty; in our task, this was the absolute difference between two brightness intensities. Note that we only used prosaccade trials, since the subcortex was shut down during antisaccade trials. Our results showed that the cortex was driving most of difficult trials and let the subcortex take over during easy trials (top panel; Fig. 4B). While the subcortical RT distribution (red; middle panel of Fig. 4B) contained little information about task difficulty, the variability among cortical RTs increased with task difficulty (blue; middle panel of Fig. 4B). The ratio between cortex- and subcortex-driven choices increased almost linearly as well (bottom panel; Fig. 4B), indicating that the model preferentially used the shallow subcortical modules for easy trials and the cortical hierarchical structure for difficult ones.

**Figure 4.**
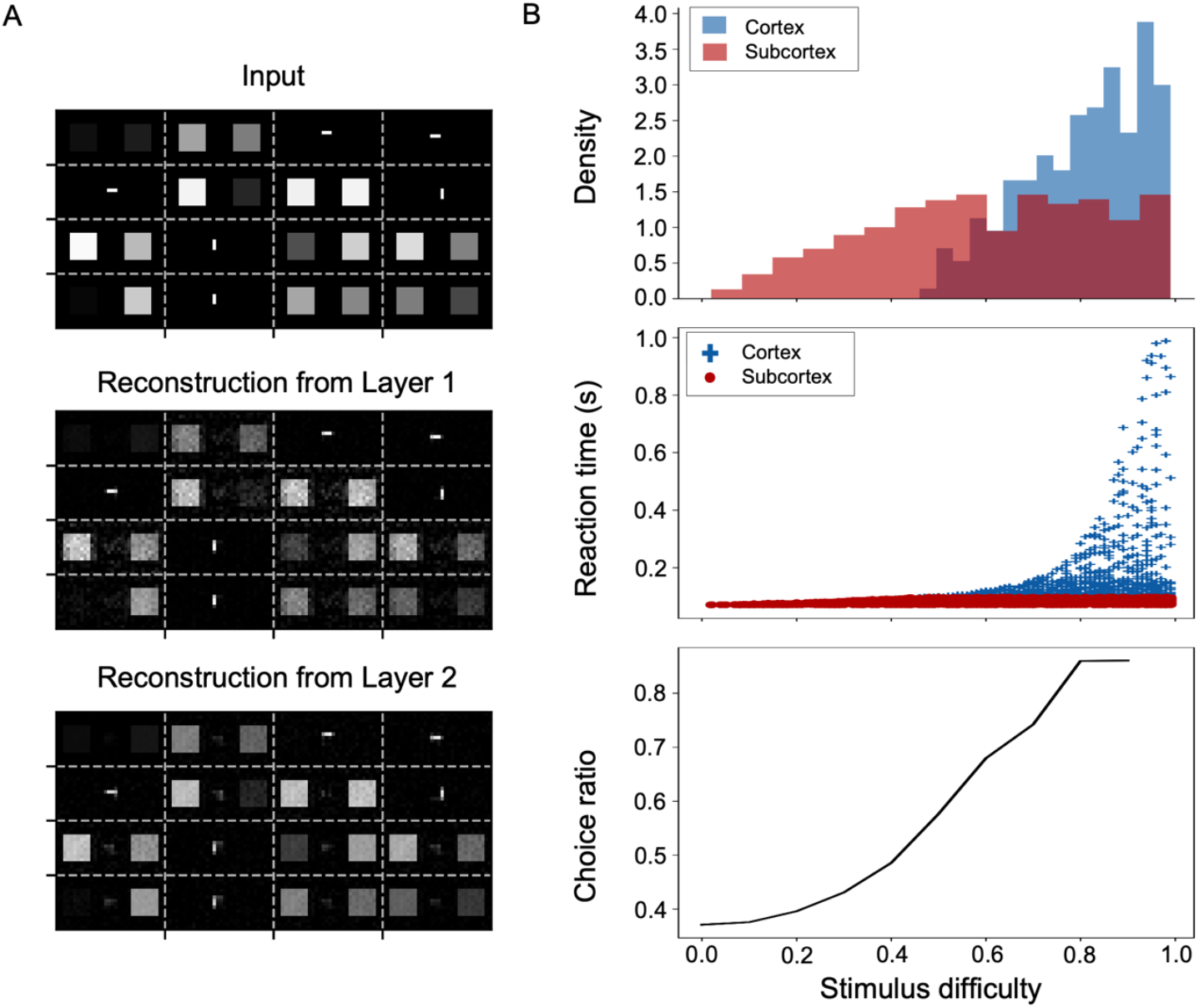
Representation learning and task difficulty in the predictive coding model. (A) Hierarchical reconstruction of sensory inputs. Sensory inputs are transmitted from the thalamus to V1 (Input layer). These inputs are then hierarchically reconstructed in the input layer from latent representations in PPC and FEF (Layers 1 and 2, respectively). Reliable reconstructions of inputs from higher layers suggest that the PC model had learned to generate useful latent representations, which can be used to perform classification and cue-based decision-making tasks. (B) Cortico-subcortical dynamics and task difficulty. During easy tasks (e.g., stimulus difficulty < 0.6), subcortical outputs primarily drive saccadic decision-making. As task difficulty increases (e.g., stimulus difficulty > 0.6), cortical outputs become more involved in the decision-making process (*top*). Subcortical reaction times, defined as latencies to reach the decision threshold, remain relatively stable across different task difficulties, while cortical reaction times exhibit increasing variability and longer latencies (*middle*). The choice ratio, i.e. the ratio of choices primarily driven by cortex over choices driven by subcortex, increases with stimulus difficulty (*bottom*). Note that the stimulus difficulty is defined as the absolute value of difference between brightness intensity values (i.e., ∣ *X*_*L*_ *− X*_*R*_ ∣).

## Discussion

Inspired by recent ideas on how shallow computational structures may underlie a significant part of brain computations^7^, we have presented a computational model that accommodates both a deep cortical hierarchy and a shallow subcortical pathway. The cortical pathway has been implemented in two ways—a convolutional feedforward network and a predictive coding network. Both versions of the cortical hierarchy can be trained to perform challenging perceptual decision tasks (in the sense of requiring rule switching, such as pro-anti, on a trial-by-trial basis); therefore, the resulting cortical pathway is adaptive and capable of learning new representations. The subcortical pathway on the other hand is shorter, simpler, and non-plastic (in the PC model), in agreement with the anatomical and physiological findings. Combining these two fundamentally different networks allows our model, and presumably real brain networks, to flexibly adapt to task demands by recruiting the most convenient structure for the job.

We decided to implement the cortical hierarchy in two ways to demonstrate the generality of our core idea, although CNN and PC structures have their own advantages and caveats. CNNs constitute a more classical way of modeling the early visual cortex, which takes inspiration from how receptive fields increase in size as information moves up the hierarchy^1,4^. This first implementation indicates that our ideas of combining CNN models with more shallow architectures fit within this standard framework, and their predictions in terms of reaction times align well with experimental observations in monkeys^13^. Lesion studies in macaques indicate the need to recruit cortical areas for the antisaccade but not for the prosaccade task^33^, which is in agreement with our model. On the other hand, PC structures work with a different framework of visual perception, which views the brain as an inference machine that generates predictions which are to be matched by expected stimuli^25–28,34^. In this sense, a mismatch signals a deviation from the internal world model which has to be accounted for in the future and triggers learning. Despite relying on a fundamentally different assumption, our model is also able to work within a PC framework for modelling the cortical hierarchy, generating robust internal representations of known stimuli and mimicking the reaction time patterns observed experimentally. Interestingly, our model self-organizes to use the cortical hierarchy or subcortical shallow network to lead the task decision depending on the difficulty of the task (Fig. 4B, bottom panel), with the subcortex leading decisions on easy trials and cortex leading on more difficult ones. This again agrees with previous results on the function of cortical and subcortical areas as a function of task complexity^22,33^.

Our results highlight the importance of considering subcortical structures when building models of sensorimotor tasks. Previous computational models have shed light on how communication between cortical circuits and subcortical areas, including thalamic nuclei^35,36^, basal ganglia^37,38^ and cerebellum^39–41^ actively shape neural dynamics during perceptual and cognitive tasks, and other structures like the brainstem might play an important role within the shallow brain architecture too^7^. Recent work, in particular, has revealed how predictive coding mechanisms may benefit from incorporating subcortical areas as ‘gist*i* pathways^42^, also in more realistic theoretical frameworks including spiking neural networks^43^. This opens the door to seriously consider the advantages of deep-shallow parallel pathways for predictive coding, taking advantage of recent advances in PC modeling^44,45^. Likewise, shallow brain architectures could be relevant for cognitive tasks in which short reaction times and hierarchically-distributed computations across cortex coexist, such as working memory^46–48^, categorization or decision making^49^. Future studies should determine how these structures contribute to perceptual tasks as well as to brain function as a whole.

## Methods

### Behavioral task

To simulate context-dependent pro- and antisaccade responses, we adapted a saccade task used for monkeys^50^. Instead of simply detecting a single stimulus in a peripheral location, our task required the model to compare the brightness of two stimuli (4×4 pixels per stimulus) and make a saccade choice depending on the prescribed cue (Fig. 1A). When presented with a horizontal bar (2×1 pixels), the model had to make a saccade towards a brighter square (i.e., prosaccade condition; left in Fig. 1A). On the other hand, a vertical bar (1×2 pixels) indicated an antisaccade condition, where the model had to make a saccade away from the brighter square and towards a darker square (right in Fig. 1A). The screen dimension was 16 × 32 pixels.

The training set consisted of 121 pairs, which consisted of all possible combinations of brightness values ranging between 0.0 and 1.0 with an interval of 0.1. For test sets, we increased the number of possible brightness values by decreasing the interval by ten-fold (i.e., 0.01) to introduce non-familiar brightness pairs. The training and test sets were balanced with respect to brightness, containing equal numbers of left side brighter and right-side brighter pairs.

### Neural network model and simulations

For both the feedforward and PC models, the cortical pathway module had access to the entire input space via convolving kernels (green dotted box; Fig. 1C) or dense connections (orange dotted box encapsulating the entire visual field considered; Fig. 3A). On the other hand, the subcortical pathway module consisted of two neuronal populations (*R* and *L* of the subcortical module in Fig. 1C and *S*_*R*_ and *S*_*L*_ in Fig. 3A for the feedforward and PC model, respectively), each with receptive fields covering only the region of either the left or right brightness stimulus. Hence, the subcortex could respond to the brightness comparison stimuli pairs – but not to cues, since they are indicative of the corresponding rule rather than of stimuli guiding the decision. The background noise (Fig. 2A and Fig. 3B) served as recurrent inputs from other subcortical neuronal population.

To simulate neural activity in a continuous and biologically plausible manner, discrete instantaneous firing rate outputs in the feedforward model were smoothed via the following differential equation:

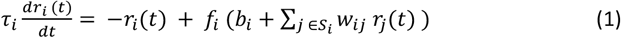

Here, τ_*i*_ is the membrane constant of neuron *i, r*_*i*_(*t*) the firing rate of neuron *i* at time t, *f*_*i*_ the activation function of the neuron (i.e., LogSoftmax for *C*_*L*_, *C*_*R*_, *S*_*L*_, *S*_*R*_, *M*_*L*_, and *M*_*R*_; ReLU for all other neurons), *b*_*i*_ the bias (i.e., intercept) of neuron *i, S*_*i*_ the set of neurons connected to the neuron *i*, and *w*_*i*._ the weight connecting neuron *j* to neuron *i*.

Since integration of synaptic inputs occurs at the neuron level, the overall layer architecture (convolutional or fully connected) does not affect Eq. 1. The temporal evolution of the firing rate is computed using the Euler method with a time step (Δ*t*) of 0.01 s:

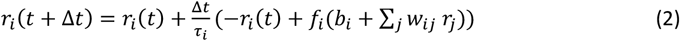

The cortical pathway of the feedforward model consisted of two convolutional (4×4) and max pooling (3×3) layers, followed by a fully connected layer with 16 neurons and 4 output neurons (L, R, P, A; Fig. 1C). It was trained on the training set via backpropagation to distinguish between pro-or anti-saccade conditions during cue presentation and to identify the side with a brighter square (left or right) during stimulus presentation. To implement cue-dependent switching in decision strategy, the final cortical output (*C*_*L*_ or *C*_*R*_) integrated the sequential classification results of the cue and stimuli in the following two steps: 1) the classification result of cue was saved in memory (not shown in the figure) and retrieved during the subsequent stimulus presentation, which in neurobiological terms may be supported by working memory; and 2) second, classification results of cue and stimulus were combined via a hard-wired hidden layer (the second array of grey squares) that acted as an XOR gate (XOR).

The subcortical pathway consisted of an input layer with two neurons, each receiving the brightness value of the left and right square (*R* and *L* of the subcortical module in Fig. 1C), a hidden layer with 16 neurons (grey squares in the subcortex module), and two output neurons (*S*_*L*_ and *S*_*R*_), indicating the subcortical decision. It was trained via backpropagation to identify the side with a brighter square (left or right) during stimulus presentation.

The three layers in the cortical module of PC model were modelled and trained according to the Rao *S* Ballard implementation^26^ (Fig. 3A). Each layer consisted of 512, 144, and 16 neurons, respectively. These layers were trained on the training set to minimize prediction errors, thereby learning to generate latent representations of inputs (i.e., both cues and stimulus). Then, a linear classifier was trained on the latent representations of the last layer to distinguish between pro- and anti-saccade conditions given a cue and to identify the side with a brighter square given a brightness comparison stimuli pair. The same two steps taken for the FF model were used to implement cue-dependent switching in decision strategy: integration of sequential classification results of cue and stimulus via a hard-wired XOR gate. However, instead of using an external memory to integrate sequential classification results of cue and stimulus, the two cue classifier neurons (P and A) had excitatory, self-recurrent connections. Once the network classified a cue, the corresponding neuron sustained its activity before and during the subsequent decision phase on a brightness comparison stimulus pair and influenced the network to make a cue-based decision.

A simplified version of a previously described recurrent network for perceptual decision-making^32^ was implemented as the subcortical module. Two excitatory neuron populations (S_L_ and S_R_; Fig. 3A), each containing 128 neurons, interacted with an inhibitory neuron population (I_S_) of 32 neurons (S_I_). The interaction between these three subcortical neuron populations were governed by two excitatory (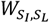 and 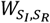), two inhibitory (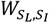 and 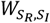), and two self-recurrent (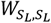 and 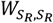) connections. The connection strengths were drawn from a uniform distribution between 0 and 1, with connection probabilities of 0.3, 0.8, and 0.4 for excitatory, inhibitory, and self-recurrent connections, respectively. The two excitatory subcortical neuron populations received inputs from the cue and stimulus via two sets of random weights (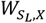 and 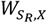) with a connection probability of 0.4. Output from these populations was then sent to motor neurons (*M*_*L*_ and *M*_*R*_) via two additional sets of random weights (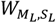 and 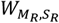) with a connection probability of 0.8.

In both the FF and PC models, the antisaccade classifier neuron (A) projected to the subcortical module, analogous to the inhibitory projections from the FEF to the SC^24^. During antisaccade trials, these projections influenced the subcortical module, which then suppressed reflexive saccades toward the brighter square, irrespective of the cued condition, thereby allowing the cortical module to mediate an antisaccade. In the FF model, the antisaccade classifier neuron made direct inhibitory projections to the subcortical decision neurons (*S*_*L*_ and *S*_*R*_). In contrast, those in the PC model indirectly suppressed the subcortical activity by projecting excitatory connections to the inhibitory neuron population (S_I_).

To evaluate the robustness of learned representations, a varying level of Gaussian noise (ϵ ~ N(0.0, σ) where σ = {0.0, 0.01, …, 0.2}) was added to both the cues and brightness stimuli. Both the feedforward and PC models were tested 100 times on the same, balanced 1000 pairs of brightness comparison stimuli for statistical comparison.

## Acknowledgements

The authors thank Sander Bohte, Matthias Brucklacher, Giulia Moreni and Parva Alavian for useful discussions.

## Author contributions

KL, MS and JFM conceived and designed the study; KL and LB performed the research; KL, LB, MS and JFM analyzed the results; KL, MS, CMAP and JFM wrote the manuscript.

## Data and materials availability

All information needed to reproduce the results of this manuscript are in the main text and Methods section. The code used to generate the results will be made openly available upon publication of this work.

